# Temporal dynamics of fluorescence and photoacoustic signals of a Cetuximab-IRDye800 conjugate in EGFR-overexpressing tumors

**DOI:** 10.1101/2024.11.26.625469

**Authors:** Mohammad A. Saad, Derek Allen, Allison Sweeney, Marvin Xavierselvan, Srivalleesha Mallidi, Tayyaba Hasan

## Abstract

Molecular fluorescence-guided surgery has shown promise for tumor margin delineation but is limited by its depth profiling capability. Interestingly, most fluorophores, either clinically approved or in clinical trials, can also be used as photoacoustic contrast agents, yet their use is limited due to the low light fluence permitted for clinical use and the limited sensitivity of current photoacoustic imaging systems. There is therefore an urgent unmet need to establish methods for enhancing contrast in molecular targeted PA imaging which could potentially complement and overcome limitations in molecular fluorescence guided therapies. In this study, we compare the photoacoustic (PA) and fluorescence imaging capabilities of a cetuximab-IRDye800 conjugate in a subcutaneous tumor xenograft model. We demonstrate that while the fluorescence signal increases steadily over time after administration of cetuximab-IRDye800, PA signal peaks early (∼2 fold higher at 6-hour as compared to pre-injection controls) and then decreases (∼1.3 fold higher at 24-hour as compared to pre-injection controls). This pattern aligns with previous findings using other antibody-conjugated PA contrast agents. Mechanistically, we demonstrate that the formation of H-aggregates upon antibody conjugation enhances PA contrast of the IRDye800. The disruption of these H-aggregates, as the antibody-dye conjugate is degraded post receptor-mediated endocytosis, decreases PA signal intensity. The timeframe of maximum PA signal and decrease thereafter is consistent with the time frame of receptor-mediated endocytosis of cetuximab-IRDye800. Our data suggests that tumor cell surface binding results in peak PA signal while lysosomal localization and degradation results in a significant drop in PA signal. Our study sheds light on the distinct temporal dynamics of PA and fluorescence signals of Cetuximab-IRDye800 conjugate and we propose that optimizing IRDye800 conjugation to antibodies can further enhance PA signal intensity when timed to precisely to capture IRDye800 in an H-aggregate form.

## Introduction

Fluorescence imaging, either through monitoring inherent variations in autofluorescence of diseased and healthy tissues or through externally administered fluorescent contrast agents, has been the imaging of choice for delineating tumor margins during surgical resection [1]. This is exemplified by the use of several fluorescent contrast agents (methylene blue, 5-Aminolevulinic acid and Indocyanine green (ICG)) for imaging sentinel lymph nodes, vascular/lymphatic pathologies and tumor tissues during surgical procedures [2-4]. However, fluorescence imaging of externally administered contrast agents is limited by superficial imaging depths of 1-2 mm as well as their non-specific accumulation leading to inaccurate margin definition. In this context, conjugating fluorophores to molecular targeted agents has been shown to improve specificity [5-8], and several clinical studies are being conducted to study the efficacy of molecular targeted fluorophores in head and neck cancer surgery (NCT01987375), pancreatic cancer (NCT02736578), prostate cancer (NCT02048150), breast cancer (NCT01508572), and rectal cancer (NCT01972373).

In addition, complementing fluorescence with photoacoustic imaging provides the much-needed depth profiling to enhance spatial resolution [9-11]. Photoacoustic (PA) imaging relies on excitation of chromophores using nanosecond pulsed lasers which induce transient thermal expansions that can be detected using ultrasound transducers [12]. In essence, PA imaging combines optical and ultrasound imaging often proving advantageous due to the combined benefits of the two imaging modalities – sensitivity of optical imaging with the imaging depth of ultrasound imaging. PA imaging is in clinical trials to evaluate its diagnostic efficacy for several types of cancer including breast cancer (NCT03897270), thyroid cancer (NCT04248166), colon and rectal tumor tissue (NCT04339374) and head and neck cancer (NCT04428515) amongst others. However, contrast free photoacoustic imaging has limited sensitivity to molecular changes as well as reduced specificity in delineating tumor from background tissues. Several contrast agents have been proposed for PA imaging, but they lack tumor specificity due to non-specific accumulation. More recently, molecular-targeted contrast agents, reported primarily for fluorescence imaging, have been studied for PA imaging in pre-clinical tumor models and for ex vivo identification of lymph node metastasis [13-16].

In the present study, we compare the PA and fluorescence imaging capabilities of a cetuximab-IRDye800 conjugate in a subcutaneous tumor xenograft model. Cetuximab is a clinically approved anti-EGFR targeted antibody and Cetuximab-IRDye800 is in clinical trials for fluorescence image guided surgery of head and neck cancers. EGFR is a promising molecular marker which has been used as a targeting molecule for imaging and therapeutic purposes due to its over-expression in several cancer types including head and neck cancers, pancreatic cancers, ovarian cancers, etc., [9, 11, 16]. Several NIR-dyes, currently used as fluorophores have also been used as contrast agents for photoacoustic measurements. Fluorescence properties of dyes depend on the radiative transfer of energy from the excited singlet state (S_1_) to the ground state (S_0_). IN contrast, photoacoustic properties depend on the non-radiative transfer of energy either from the excited singlet state (S_1_) or the excited triplet state (T_1_), achieved through an inter-system crossing. In principle, NIR-dyes with high fluorescence quantum yields (Φ_FL_) are often preferred for fluorescence-based imaging, while dyes, characterized by a relatively higher non-radiative quantum yield (Φ_NR_, quantified as 1 - Φ_FL_) can be used as photoacoustic contrast agents. Of the several fluorescent dyes in clinical use ICG (Φ_FL_ = 0.12) and IRDye800CW (Φ_FL_ = 0.15) [17], are relatively better photoacoustic contrast agents and have been explored for PA imaging. The imaging contrast of these agents can be significantly altered depending on their aggregation status, concentration, environment and protein-binding [14, 18, 19]. In the present study, using a Cetuximab-IRDye800 conjugate we report that conjugation of IRDye800 to an antibody facilitates its H-aggregation thereby enhancing PA signal intensity. However, with the tendency of cell surface receptor-targeted antibodies to undergo endocytosis, when applied to in vivo tumor imaging the PA contrast is reduced as the Cet-IRDye800 conjugate is internalized and degraded in the lysosomes. As conventional wisdom holds that prolonged incubation times (2-5 days in clinical studies), maximize the signal to background ratio in fluorescence imaging using antibody dye conjugates, we demonstrate that this may negatively influence PA contrast. Through comparative microscopic tumor distribution studies, we show that although the distribution of Cetuximab-IRDye800 in tumor tissue sections is low at early timepoints it, however, is specific to the tumor tissues. For these reasons, implementing Cetuximab-IRDye800 based PA imaging at early timepoints post-infusion may be ideal for diagnostic applications such as sentinel lymph node imaging to identify occult metastases.

## Materials and Methods

### Synthesis of Cetuximab-IRDye800-AF647 (Cet-IRDye-AF) conjugate

The human anti-EGFR antibody (Cetuximab, Erbitux from Eli Lilly) was obtained from Premium Health Services (Columbia, MD, USA). Before initiating the conjugation reaction, the antibody was filtered by passing through a 0.22 µm syringe filter followed by estimating the concentration using a Pierce BCA protein assay kit (ThermoFisher Scientific, Waltham, MA, USA). The conjugation reaction was initiated by adding N-hydroxysuccinimide (NHS) esters of IRDye800 (LI-COR Biosciences, Lincoln, NE, USA) and AF647 (ThermoFisher Scientific, Waltham, MA, USA) at a molar excess of 4 and 2, respectively, to 2 ml of ∼ 2 mg/ml Cetuximab. The molar concentration of IRDye800 and AF647 was varied to synthesize Cet-IRDye-AF with varying degree of labelling for the two dyes. The reaction was allowed to proceed for 4 h under continuous stirring at room temperature followed by purification using a 7 kDa MWCO Zeba Spin Desalting Column (ThermoFisher Scientific, Waltham, MA, USA) pre-equilibrated with PBS. The purified Cetuximab-IRDye800-AF647 (Cet-IRDye-AF) conjugate was then concentrated using a 30 KDa Amicon Ultra-15 Centrifugal Filter Unit (MilliporeSigma, Burlington, MA, USA). The final purified Cet-IRDye-AF conjugate was stored at 4 °C.

### Characterization of Cet-IRDye-AF conjugate

The degree of labelling of IRDye800 on Cetuximab was calculated by monitoring the absorption of Cet-IRDye-AF conjugate, using a Thermo Evolution 300 – UV-Vis spectrophotometer (ThermoFisher Scientific, Waltham, MA, USA), at 780 nm using the molar extinction coefficient of 270,000 M^−1^ cm^−1^ for IRDye800 at 780 nm in 1:1 PBS: methanol. AF647 concentration was calculated using the molar extinction coefficient of 239,000 M^−1^ cm^−1^ at 650 nm in PBS with an 11% correction factor for IRDye800 at 650 nm. Cetuximab concentration was calculated using a Pierce BCA protein assay kit (ThermoFisher Scientific, Waltham, MA, USA).

SDS-PAGE analysis was performed by mixing the Cet-IRDye-AF conjugate with 1X Laemmli buffer (Boston BioProducts (Milford, MA, USA), heating at 95 °C for 5 min and loading it on a 10% precast polyacrylamide gel (Bio-Rad, Hercules, California, USA). The separated bands were visualized using a ChemiDoc MP Imaging System (Bio-Rad, Hercules, California, USA). Image analysis and quantification were performed using ImageJ (National Institute of Health).

Comparative fluorescence studies for free AF647, IRDye, and Cet-IRDye-AF were performed by diluting the stock of the dye or conjugate in PBS and mixing it with 1.2% agarose in a 1:1 ratio, followed by overlaying it on an agarose (1.2%, 100 µl) bed in a well of a 96-well plate. The gels were allowed to solidify and imaged in hyperspectral mode using an IVIS imaging system using an IVIS Lumina Series III In Vivo Imaging System (PerkinElmer, Waltham, MA, USA). Baseline spectra of free AF647 and IRDye800 phantoms were also measured and used for unmixing and generating unmixed composite images of IRDye800 and AF647 in the Cet-IRDye-AF phantoms. Quantification of fluorescence intensities was performed using the Living Image software provided by the manufacturer.

Photoacoustic imaging (PAI) was performed using a Visualsonics Vevo2100 LAZR system (Visualsonics, Fujifilm, ON, Canada). The 96-well plate with the phantoms was submerged in double-distilled water and the PA transducer (LZ220), with bubble-free ultrasound gel applied to it, was placed in close proximity to the sample containing phantoms. Image analysis was performed using the built-in software provided by Visualsonics, Fujifilm (ON, Canada). Imaging parameters, laser power, photoacoustic gain, persistence, and frame rate, were kept constant across different measurements.

For trypsinization studies, Cet-IRDye-AF at a concentration of 20 µM IRDye800 equivalent was mixed with 1% trypsin and incubated at 37 °C was 24 h. Following this, phantoms were prepared and imaged as described previously.

### Generation of subcutaneous A431 xenograft tumors

A431 cells cultured in DMEM (Corning, Corning, NY, USA), supplemented with 10% FBS, and a mixture of penicillin (50 international units (I.U.)/mL and streptomycin (50 µg/mL) (Corning, Corning, NY, USA). All animal studies were performed in accordance with protocols approved by the Institutional Animal Care and Use Committee (IACUC). For tumor implantation, confluent cell cultures were trypsinized, counted and resuspended in DMEM. 2.5 x 10^6^ cells in 50 µl of 50% matrigel were implanted subcutaneously in 6- to 8- week-old athymic nude mice (The Jackson Laboratory, Bar Harbor, ME, USA). After two weeks the animals were imaged for background fluorescence and photoacoustic signal followed by intravenous injection of Cet-IRDye-AF (250 µg Cetuximab equivalent) through the lateral tail vein. The animals were imaged again at 6 h, 12 h and 24 h. A cohort of animals was sacrificed at 6 h and 24 h each, respectively, for histopathological evaluation of tumor tissues.

In vivo fluorescence measurements were carried out on isoflurane anesthetized mice in hyperspectral mode using an IVIS Lumina Series III In Vivo Imaging System (PerkinElmer, Waltham, MA, USA). Baseline spectra of free AF647 and IRDye800 phantoms were used for unmixing and generating unmixed images of IRDye800 and AF647. Quantification of fluorescence intensities was performed using the Living Image software provided by the manufacturer.

Photoacoustic imaging (PAI) was performed using a Visualsonics Vevo2100 LAZR system (Visualsonics, Fujifilm, ON, Canada). Mice were anesthetized and placed in prone position with bubble-free ultrasound gel applied on the tumors. The PA transducer (LZ220) was placed in proximity to the tumor and images were acquired either in a single plane, multiwavelength mode or in 3D mode at pre-defined wavelengths to differentiate between endogenous hemoglobin (Hb) signals and IRDye800 signals. Co-registered B-mode images were acquired for identifying tumor planes between different imaging time-points. Imaging parameters, laser power, photoacoustic gain, persistence, and frame rate, were kept constant across different measurements. Raw data were exported as ‘.xml’ files from the Vevo LAB software and then imported to MATLAB. The hemoglobin and dye content were unmixed from the multi-wavelength data using the linear least squares method [20] via the lsqnonneg MATLAB function. The molar extinction coefficients for oxygenated and deoxygenated Hb were gathered from S.A. Prahl [21] while the IRDye800 dye molar extinction coefficients were taken directly from the spectrophotometer readings described in the section above. Images were thresholded for display purposes and shown in 2D and 3D with the same dynamic range. Quantification was performed across the whole tumor volume and taken as a pixel-wise average for IRDye800 and HbT amplitude values.

### Immunofluorescence Staining and Image Analysis

Cet-IRDye-AF injected mice were sacrificed in accordance with the IACUC approved procedures. Tumor tissues were collected and embedded in Tissue-Tek O.C.T. compound (Sakura, Torrance, CA, USA) and frozen until further use. Tumor tissues were then cryosectioned at a thickness of 5 μm, fixed with acetone, stained using primary anti-EGFR (EP38Y; Abcam, Cambridge, United Kingdom) and anti-CD31 (MEC 13.3; Woburn, MA, USA) antibodies followed by secondary Goat anti-Rabbit IgG (H+L) Highly Cross-Adsorbed Secondary Antibody, Alexa Fluor™ Plus 488 (ThermoFisher Scientific, Waltham, MA, USA) and Goat anti-Rat IgG (H+L) Cross-Adsorbed Secondary Antibody, Alexa Fluor™ 546 (ThermoFisher Scientific, Waltham, MA, USA) antibodies. Rabbit IgG, monoclonal (EPR25A, Abcam, Cambridge, United Kingdom) and Rat IgG2a kappa Isotype Control (eBR2a, eBioscience, ThermoFisher Scientific, Waltham, MA, USA) were used as isotype control. The sections were counterstained with ProLong™ Glass Antifade Mountant with NucBlue™ Stain (Invitrogen, ThermoFisher Scientific, Waltham, MA, USA) and imaged using a Hamamatsu NanoZoomer slide scanner (Hamamatsu Photonics, Shizuoka, Japan). Image acquisition settings were consistent for each tumor section and images were acquired for 4 channels which contained DAPI, EGFR (FITC), CD31 (TRITC), as well as the endogenously injected free and Cet-IRDye-AF-bound Alexa Fluor 647. Images were separated into their constituent channels using the NDP.View2 application and subsequently loaded into MATLAB. Control mouse sections without injected Cet-IRDye-AF were used to determine threshold values for true Cet-IRDye-AF signal. For EGFR and CD31 signal, isotype antibodies were used for threshold setting. For each slide, the tumor, skin, and muscle tissue sections were identified in accordance with H&E, EGFR, and DAPI stains. After the different tissue types were identified, the thresholds were applied, and the images were cropped based on the specific tissue type of interest. Mean Cet-IRDye-AF intensity and percent area were determined via image matrix calculations. For determining the separation between components of different fluorescent channels, the pdist2 MATLAB function was used.

### Fluorescence and photoacoustic imaging of in vitro tumor phantoms

Tumor phantoms were prepared as described in our previous studies [22-24]. Briefly, A431 cells were seeded in 150 mm dishes and treated with Cet-IRDye-AF for the indicated time points following which cells were collected by gently scraping by a cell scraper, counted and resuspended in 5% gelatin. The resuspended cells (1 x 10^7^) were immediately poured in a well of a 96-well plate previously prelaid with 10% gelatin (100 μL) to minimize reverberations from the base of the 96-well plate during PA imaging. The tumor phantoms were allowed to solidify and imaged within 24 h. IVIS and PAI were performed as described earlier.

### Confocal imaging for assessing Cet-IRDye-AF localization

A431 cells were seeded onto glass bottom 35 mm cell culture dishes at a density of 300,000 cells per dish. Cet-IRDye-AF was added at an AF647 equivalent concentration of 250 nM. 30 min prior to the end of incubation, cells were incubated with LysoTracker® Green DND-26 (Cell Signaling Technology, Danvers, MA, USA) at a concentration of 50 nM, fixed with 4% paraformaldehyde, counterstained with Hoechst (ThermoFisher Scientific, Waltham, MA, USA) and visualized under a confocal microscope (Olympus FV1000, Shinjuku City, Tokyo, Japan). Quantitative analysis of colocalization were performed after converting images to 8-bit and subtracting background using a rolling ball radius of 50 through ImageJ (National Institute of Health).

### SDS-PAGE analysis

A431 cells were seeded onto 35 mm cell culture dishes at a density of 300,000 cells per dish. Cet-IRDye-AF was incubated at an AF647 equivalent concentration of 250 nM, and the cells were collected by gently scraping by a cell scraper at the end of the desired incubation period and pelleted. Cell pellets were lysed using RIPA lysis buffer and protein estimation in the cell lysate was performed using a Pierce BCA protein assay kit (ThermoFisher Scientific, Waltham, MA, USA). 20 µg of cell lysate was then mixed with 1X Laemmli buffer (Boston BioProducts (Milford, MA, USA), heated at 95 °C for 5 min followed by loading on a 10% precast polyacrylamide gel (Bio-Rad, Hercules, California, USA). The separated bands were visualized using a ChemiDoc MP Imaging System (Bio-Rad, Hercules, California, USA). Image analysis and quantification were performed using ImageJ (National Institute of Health).

## Results and Discussion

### Cet-IRDye-AF synthesis

Cetuximab-IRDye800-AF647 (Cet-IRDye-AF) synthesis was performed by a previously established protocol [22, 25, 26] with slight modifications as described in **Fig 1A**. The FDA approved anti-EGFR antibody, Cetuximab, was used for specificity and targeting [27], and the synthesis protocol was optimized to incorporate Alexa Fluor 647 for ex vivo fluorescence imaging in tumor sections. The degree of labeling (DOL) was 1.87 ± 0.17 and 0.87 ± 0.17 for IRDye800 and AF647, achieved using dye to antibody ratios of 4 and 2, respectively (**Fig 1D**). As shown in **Fig 1B**, Cet-IRDye-AF showed absorption peaks corresponding to both AF647 and IRDye800. There was a spectral overlap between the two dyes where IRDye800 contributed to the AF647 absorption and a correction factor of 0.11 was applied for calculating AF647 concentrations. The interference of AF647 absorption with that of IRDye800 at 780 nm was minimal. The purity of the Cet-IRDye-AF conjugate was confirmed by SDS-PAGE (**Fig 1C**), and as shown both dyes were bound to the heavy and light chains of the antibody with minimal (< 2%) free dye in the solution. Increasing the IRDye800 to Cetuximab ratio in the reaction mix led to a drastic decrease in the conjugation efficiency of AF647. The conjugation efficiency of IRDye800 however was not affected with increasing IRDye800 to Cetuximab ratio and showed a slight decrease (**Fig 1D**).

**Figure 1:**
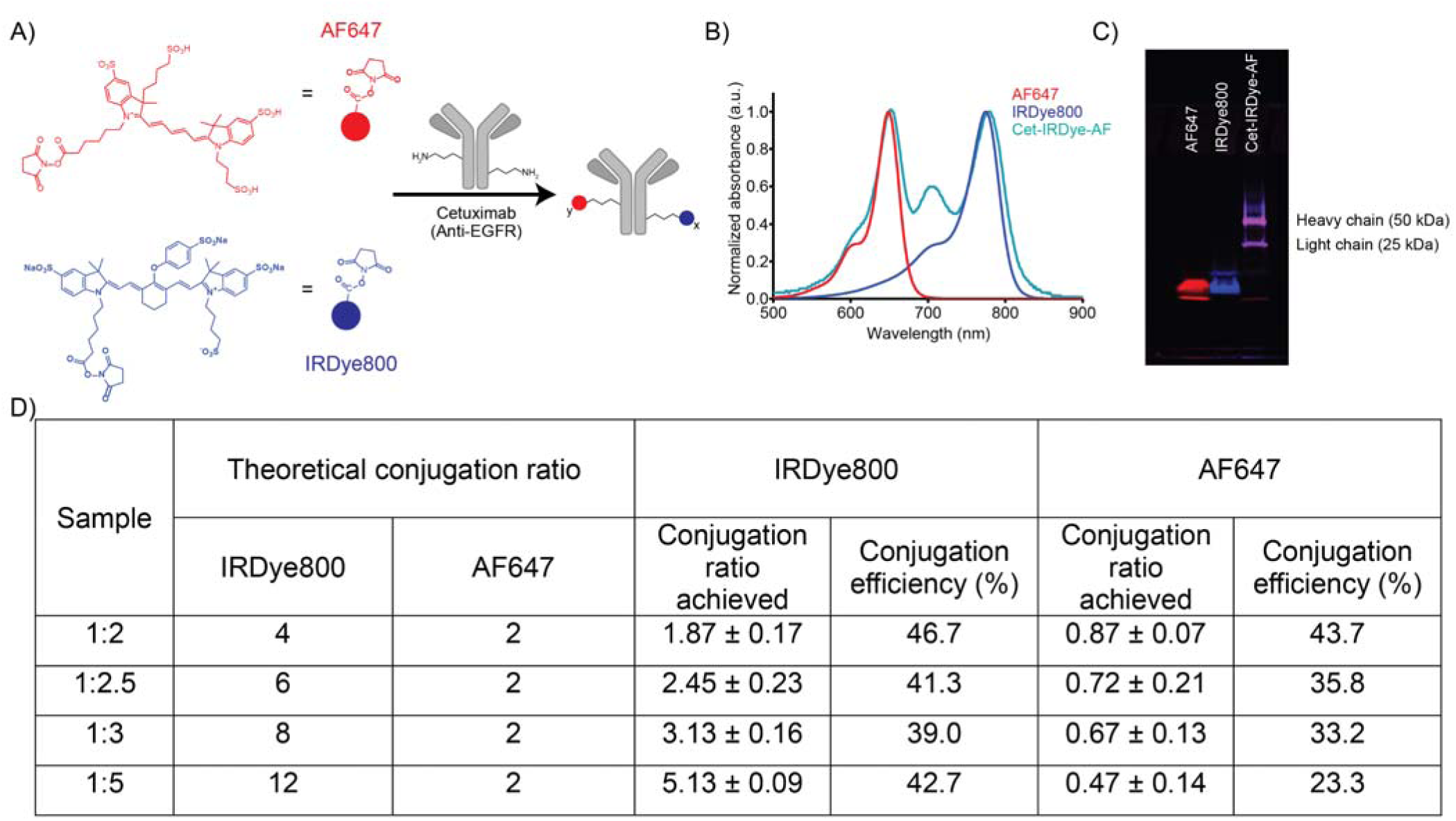
Synthesis and characterization of Cet-IRDye-AF conjugate. **(A)** Schematic representation of the synthesis of Cetuximab-IRDye800-AF647 conjugate. The NHS esters of the two dyes were reacted with the anti-EGFR antibody, Cetuximab, in predefined ratios to develop the conjugate followed by purification through gel filtration chromatography. **(B)** UV-Vis absorption spectra of AF647 (red), IRDye800 (blue) and Cet-IRDye-AF (Cyan). The spectrum for Cet-IRDye-AF shows characteristic absorption peaks for both IRDye800 and AF647. **(C)** SDS-PAGE gel image of Cet-IRDye-AF showing binding of the two dyes to both the heavy and light chains of the Cetuximab antibody. Free dyes are shown in red and blue bands for reference. **(D)** Table showing the conjugation ratio of IRDye800 and AF647 per mole Cetuximab in a reaction to generate Cet-IRDye-AF with varying IRDye800 ratios, as mentioned in the sample name. The conjugation ratios for IRDye800 and AF647 are listed along with the conjugation efficiency. Data are presented as mean ± SD. n ≥ 2 per sample.

### Distinct temporal dynamics of PA and fluorescence signals of Cet-IRDye-AF

Tumors were developed by subcutaneous implantation of A431 cells in the flank of Swiss nu/nu mice. After 2 weeks, tumor bearing mice were imaged to record the background fluorescence and PA signal followed by intravenous administration of Cet-IRDye-AF. The treated mice were then imaged at 6 and 24 hours, post-injection (**Fig 2A**). As shown in **Fig 2B**, mice treated with Cet-IRDye-AF showed an increase in IRDye800 fluorescence post-injection. Quantification of fluorescence images revealed a significant increase in fluorescence signal, 6 h post Cet-IRDye-AF injection (∼4000-fold over pre-injection levels, p = 0.0039; pre-injection vs 6 h), but a non-significant increase in fluorescence beyond the 6 h time-point (∼5000-fold over pre-injection levels, p > 0.9999; 6 h vs 24 h and p = 0.0018; pre-injection vs 24 h) (**Fig 2C**). AF647 fluorescence also showed a similar trend, although the difference between fluorescence signal at 6 h and 24 h was much higher (∼1500-fold at 6 h vs ∼3000-fold at 24 h over pre-injection levels). This is possibly due to dequenching of fluorescence signal (**supplementary Fig S1 and S2**) upon cellular uptake and lysosomal degradation of the Cet-IRDye-AF conjugate (discussed later). However, the tumor to background fluorescence ratio increased significantly for both IRDye800 (p = 0.0381; 6 h vs 24 h) (**Fig 2D**) and AF647 (p = 0.0009; 6 h vs 24 h) (**supplementary Fig S1D**) with time, due to clearance of Cet-IRDye-AF conjugate from circulation and surrounding healthy tissue. This is in agreement with previous clinical and pre-clinical studies which suggest that the time between infusion and tissue sampling does not influence intratumoral uptake of antibody-dye conjugates [28], but does influence tumor to background fluorescence ratios due to clearance of the conjugates from healthy tissues.

**Figure 2:**
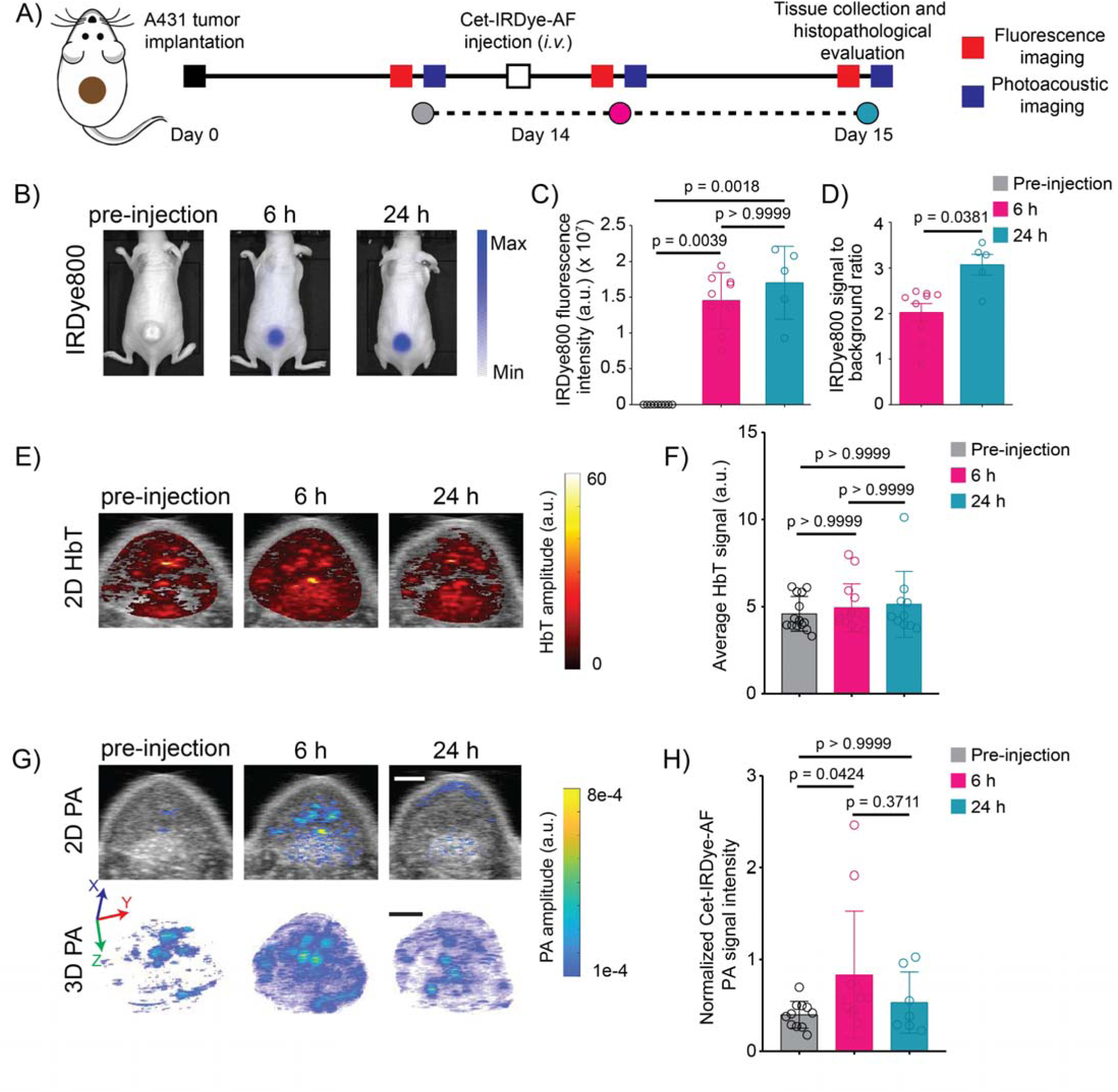
Temporal dynamics of PA and fluorescence signals of Cet-IRDye-AF. **(A)** Experimental timeline of in vivo dual modal imaging in A431 subcutaneous xenograft model. **(B)** In vivo fluorescence imaging of tumor bearing mice before injection and 6 h and 24 h after injection of Cet-IRDye-AF conjugate. **(C)** Quantification of absolute IRDye800 fluorescence signal intensity at different time-points before and after Cet-IRDye-AF injection. **(D)** Tumor to background fluorescence ratio at different time-points before and after Cet-IRDye-AF injection. **(E and F)** 2D HbT images and their quantification at the indicated timepoints. **(G and H)** 2D and 3D render of IRDye800 PA signal amplitude in the tumor, post spectral unmixing, overlaid on ultrasound image shown in grayscale and its quantification at the indicated timepoints (scale bar: 2 mm). Data are presented as mean ± SD (n ≥ 5 per group), analyzed using Kruskal-Wallis test, followed by post-hoc pairwise comparisons using Dunn’s test. P-values are provided for each graph.

In parallel to fluorescence imaging, we performed spectroscopic photoacoustic imaging (**Fig 2A**) on these tumor bearing mice. As shown in **Fig 2E and 2F**, the total Hemoglobin (HbT) signal across different time-points, pre- and post-Cet-IRDye-AF administration, was similar suggesting minimal variations in vasculature, perfusion and Hb levels. However, the PA signal of IRDye800 increased at 6 h post-Cet-IRDye-AF injection (∼2-fold over pre-injection levels, p = 0.0424) followed by a slight decrease (∼1.3 fold over pre-injection levels, p > 0.9999; and p = 0.3711 for 6 h vs 24 h) by the 24 h time point (**Fig G and H**). A similar trend of an early peak in PA signal intensity has been reported previously with a different molecular targeted dual imaging probe (Trastuzumab BHQ3 and AF488) [9]. Based on the temporal discrepancy between the fluorescence and PA signal maxima, we hypothesized that the decrease in PA signal from 6 h to the 24 h time-point may be due to variations in microscopic tumor localization, rather than macroscopic tumor accumulation of the Cet-IRDye-AF conjugate.

### Variations in intra-tumoral distribution of Cet-IRDye-AF with time

To assess the intratumoral localization of Cet-IRDye-AF, tumor tissues from Cet-IRDye-AF injected mice were harvested at either 6 h or 24 h, post-injection, and evaluated for microscopic distribution of Cet-IRDye-AF conjugate. Uninjected mice tumors were used as controls for thresholding fluorescence signals. At both 6 h and 24 h post Cet-IRDye-AF administration, tumor fluorescence signal was significantly higher as compared to the fluorescence signal in the overlying skin/dermal tissue and surrounding muscle tissue (**supplementary Fig 2**), suggesting specificity of the Cet-IRDye-AF conjugate, even at an early timepoint. At 24 h post-injection, the tumor fluorescence intensity per pixel showed an increase, although not significant, as compared to 6 h (**Fig 3C**; p = 0.4661 for 6 h vs 24 h). In addition, the percentage tumor coverage of Cet-IRDye-AF was higher (**Fig 3D**), although not significant, suggesting higher distribution of the conjugate at 24 h post-injection (p = 0.1124 for 6 h vs 24 h). We then calculated Cet-IRDye-AF intensity normalized for vasculature (Cet-IRDye-AF to CD31 ratio) to account for variations in the vasculature in the different sections. Similar to the previous observations on Cet-IRDye-AF intensity in the tumor and its percentage tumor coverage, the Cet-IRDye-AF to CD31 ratio showed an increase (not significant, p = 0.0767 for 6 h vs 24 h) at 24 h as compared to the 6 h time point **(Fig 3E)**. We also calculated the Cet-IRDye-AF distance from the nearest blood vessel (CD31) as a metric to quantify distribution. The Cet-IRDye-AF distance from the nearest blood vessel was significantly higher in the 24 h group as compared to the 6 h group (p = 0.0439) **(Fig 3F).** This suggests that at the 24 h timepoint slightly more Cet-IRDye-AF can enter the tumor and extravasate into deeper tumor layers farther away from the blood vessels. The Cet-IRDye-AF to EGFR ratio, a parameter used to determine the number of available EGFR sites bound by the Cet-IRDye-AF, also showed a slight but non-significant increase at 24 h as compared to that at 6 h (p = 0.1434) **(Fig 3G)**.

**Figure 3:**
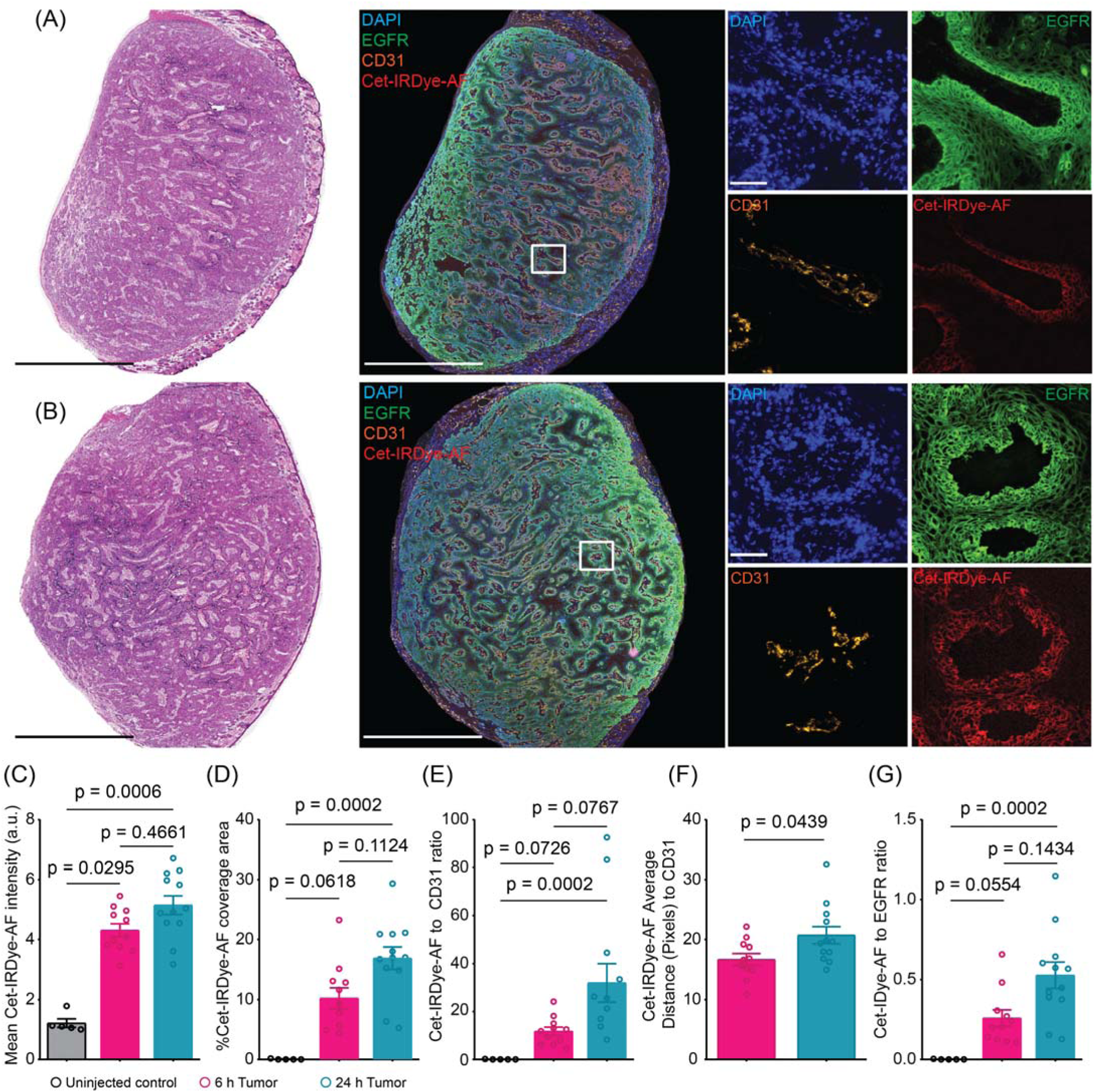
Cet-IRDye-AF distribution demonstrates variations in microscopic heterogeneity at 6 h and 24 h post-injection. **(A)** H&E and corresponding immunofluorescence image of tumor cross-section obtained from a mouse tumor 6 h after Cet-IRDye-AF injection. **(B)** H&E and corresponding immunofluorescence image of tumor cross-section obtained from a mouse tumor 24 h after Cet-IRDye-AF injection (Scale bar: 2.5 mm, inset scale bar: 100 µm). **(C-G)** Quantification of Cet-IRDye-AF distribution heterogeneity parameters including mean Cet-IRDye-AF intensity, %Cet-IRDye-AF coverage area **(D)**, Cet-IRDye-AF to CD31 ratio to quantify the amount of Cet-IRDye-AF per blood vessel **(E),** distance of Cet-IRDye-AF from nearest CD31+ blood vessel **(F)**, and Cet-IRDye-AF to EGFR ratio to quantify number of EGFR bound by Cet-IRDye-AF **(G)**. Data are presented as mean ± SD (n ≥ 10 tumor sections per group from more than 5 mice each for the 6 and 24 h and 2 mice from the untreated group), analyzed using Kruskal-Wallis test, followed by post-hoc pairwise comparisons using Dunn’s test. P-values are provided for each graph.

These data suggest that a 24 h gap between infusion and tissue sampling increases microscopic in vivo fluorescence signal due to enhanced uptake and distribution of Cet-IRDye-AF conjugate. The percentage tumor coverage of Cet-IRDye-AF, Cet-IRDye-AF intensity normalized for vasculature, and Cet-IRDye-AF to EGFR ratio all showed a significantly higher value for the 24 h timepoint when compared to the pre-injected controls. These parameters although showed an increase at the 6 h timepoint, the difference was not significant. These results do support the in vivo observations of increase in macroscopic tumor fluorescence; however they failed to address the decrease in PA signal from the 6 h to the 24 h time-point.

### Variations in sub-cellular Cet-IRDye-AF localization with time

To understand the contrasting variations in IRDye800 fluorescence and PA signals, we postulated that these differences could be due to variations in the microenvironment of the IRDye800 molecules. To gain insight into Cet-IRDye-AF sub-cellular localization, Cet-IRDye-AF was incubated with A431 cells *in vitro* and imaged for localization at different time-points. As shown in **Fig 4A**, the Cet-IRDye-AF binds to the A431 cell surface within an hour followed by gradual endocytosis and localization in the lysosomes. This was validated by calculating the Pearsons colocalization coefficient which showed a gradual increase in lysosomal colocalization with incubation time. The lysosomal localization was found to be significantly higher at 24 and 48 h, post Cet-IRDye-AF incubation in comparison to that at the 1 h timepoint (p = 0.0041 for 24 h and p = 0.0002 for 48 h vs 1 h). In a separate experiment, we prepared cell lysates from Cet-IRDye-AF treated A431 cells at different time-points and subjected them to SDS-PAGE analysis (**Fig 4C**). As shown in **Fig 4C**, bands corresponding to the free dyes (AF647 and IRDye800) appear at the 12 h time-point, suggesting that at the initial time points the Cet-IRDye-AF is either accumulated on the cell surface or is in an early stage of the receptor-mediated endocytosis pathway. The relative band intensity of the free dyes increased significantly at the 24 h time-point as compared to that observed at 6 h post-Cet-IRDye-AF incubation, suggesting degradation of Cet-IRDye-AF upon lysosomal localization. These data suggest that the decrease in PA signal as observed in the *in vivo* study, may possibly be due to the lysosomal localization and degradation of the Cet-IRDye-AF conjugate.

**Figure 4:**
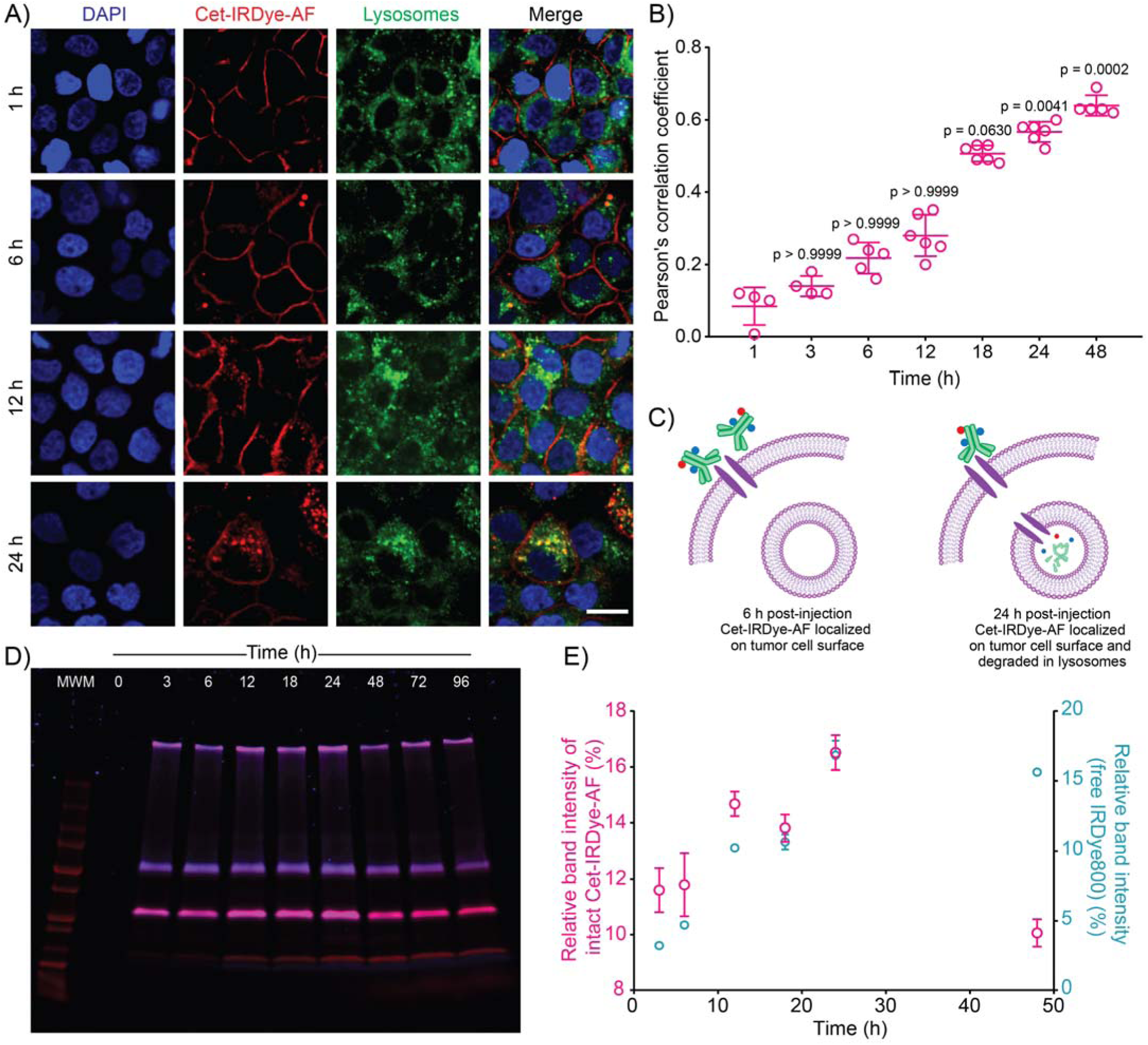
Evaluating sub-cellular localization of Cet-IRDye-AF with time in 2D A431 cultures. **(A)** Confocal images of A431 cell cultures showing localization of Cet-IRDye-AF (red) in the lysosomes (green) with time. Cell nuclei are shown in blue. (Scale bar: 20 μm) **(B)** Pearson’s correlation coefficient showing a steady increase in colocation of Cet-IRDye-AF in the lysosomes with time. Data are presented as mean ± SD (n ≥ 4 images per timepoint, analyzed using Kruskal-Wallis test, followed by post-hoc pairwise comparisons using Dunn’s test. P-values are provided for each time-point compared to the 1 h timepoint. **(C)** Pictographic representation of the status of Cet-IRDye-AF conjugate at the 6 h and 24 h timepoint. At 6 h, Cet-IRDye-AF primarily resides on the cell surface, while at the 24 h timepoint Cet-IRDye-AF is internalized and degraded in the lysosomes. **(D)** SDS-PAGE analysis of cell lysates obtained at the indicated timepoints after Cet-IRDye-AF incubation. Beyond the 6 h time point, low molecular weight bands corresponding to degraded product of the Cet-IRDye-AF conjugate were visible. **(E)** Quantification of the SDS-PAGE gels shows a significant increase in degradation product beyond the 6 h timepoint (n = 2)

### Lysosomal uptake and cleavage of Cet-IRDye-AF reduces PA signal intensity

Having established the critical time-points associated with Cet-IRDye-AF binding and cellular uptake, we performed fluorescence and PA imaging on A431 cell phantoms pre-treated with Cet-IRDye-AF for either 6, 24 or 48 hours. Cet-IRDye-AF treated A431 tumor cell phantoms were prepared by embedding 10^7^ cells in gelatin and pouring that in a well of a 96-well plate (**Fig 5A and B**). **Fig 5C and 5D** show the IRDye800 fluorescence signals from Cet-IRDye-AF treated A431 cells which, as expected, increased with increase in incubation time. A similar trend of steady increase in AF647 fluorescence signal was also observed (**supplementary Fig S3**). PA signals, in contrast, peaked early, at the 6 h time point (**Fig 5E**), and decreased thereafter (**Fig 5F**), consistent with the *in vivo* dynamics of PA signal fluctuations. Based on these findings, we proposed that the peak PA signal is achieved at early time-points when the Cet-IRDye-AF conjugate is intact and prolonging incubation results in Cet-IRDye-AF degradation in the lysosomes leading to a drop in PA signal.

**Figure 5:**
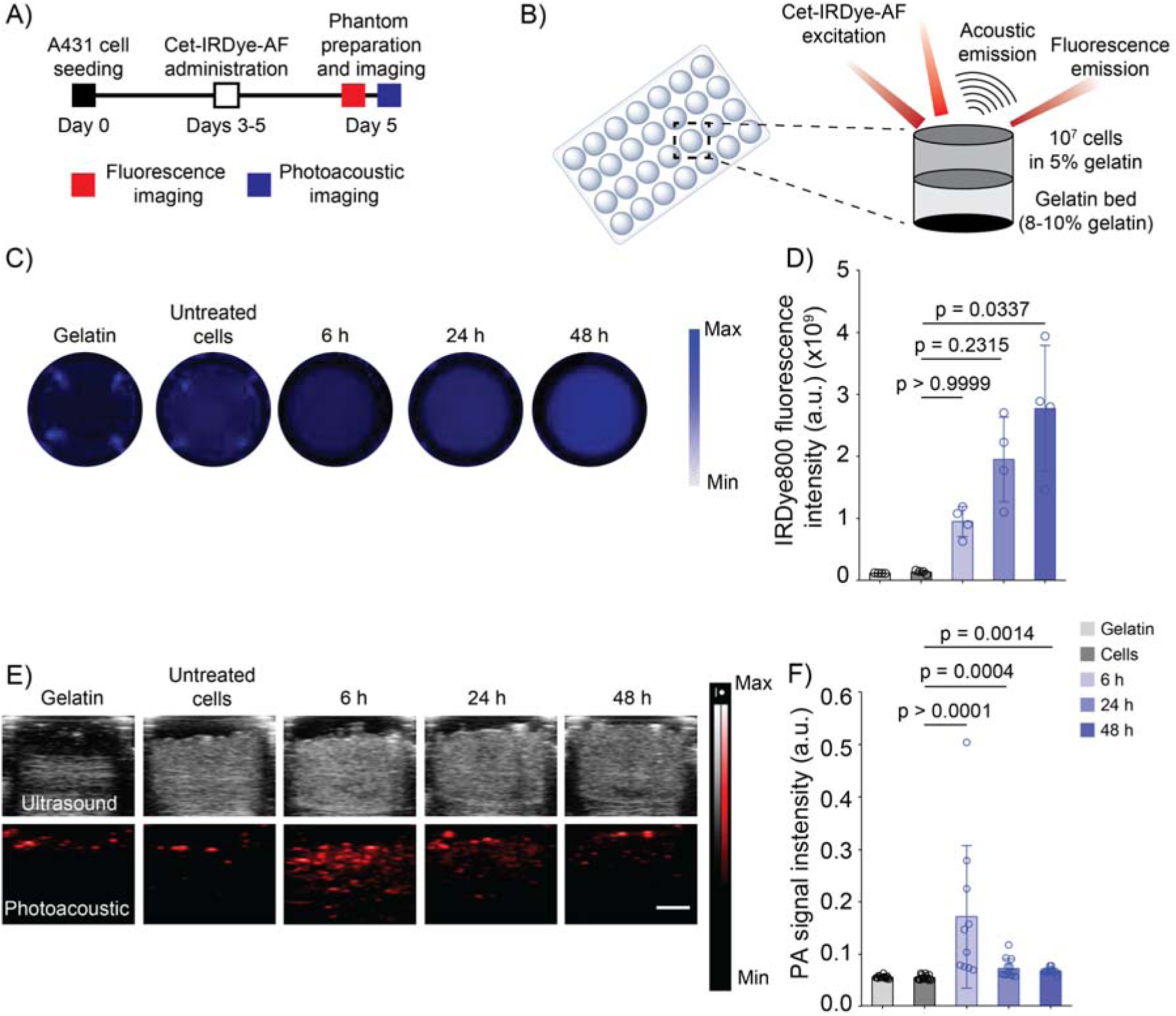
**(A)** Experimental timeline for the Cet-IRDye-AF-based in vitro multi-modal imaging. **(B)** Experimental setup for preparing phantoms for multi-modal imaging. **(C)** IVIS images of the Cet-IRDye-AF-treated A431 cells prepared as gelatin embedded tissue mimicking phantoms in a 96-well plate. **(D)** Quantification of fluorescence signals from the IVIS images. Incubation of A431 cells with Cet-IRDye-AF showed a steady increase in fluorescence signals with time. **(E)** Ultrasound and photoacoustic images of the Cet-IRDye-AF treated A431 cell phantoms. **(F)** Quantification of photoacoustic signals obtained from the photoacoustic images shows an increase in PA signal from A431 cells at 780 nm in 6 h followed by a decrease till 48 h. (Scale bar: 2 mm). Data are presented as mean ± SD (n = 4 tumor mimicking phantoms per group for fluorescence images. PA image analysis was performed on 4 tumor mimicking phantoms with 2-3 ROIs identified per phantom. Data is analyzed using Kruskal-Wallis test, followed by post-hoc pairwise comparisons using Dunn’s test. P-values are provided for each graph.

To further test our hypothesis that an intact Cet-IRDye-AF conjugate results in a higher PA signal intensity, we first recorded the fluorescence and PA signal intensity for free IRDye800 and Cet-IRDye-AF with increasing concentrations. As shown in **Fig 6A and 6C**, IRDye800 fluorescence and PA signal showed a linear increase with an increase in concentration. Interestingly, at similar concentrations Cet-IRDye800 fluorescence was significantly lower than the free IRDye800 (**Fig 6A**). However, in contrast, the PA signal from the Cet-IRDye-AF conjugate was significantly higher than the free IRDye800, suggesting that conjugation to an antibody improves PA signal amplitude (**Fig 6C**).

**Figure 6:**
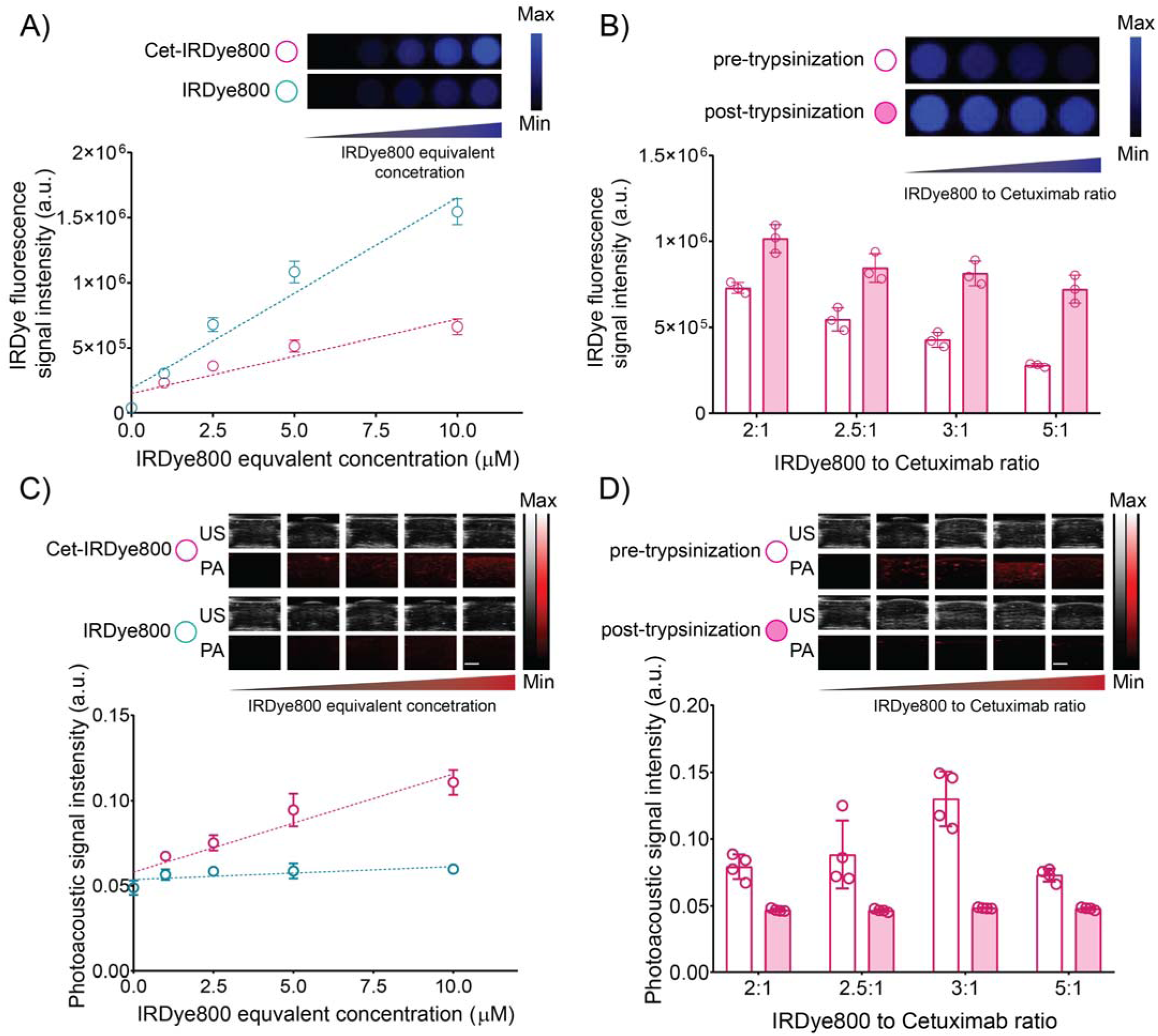
Fluorescence and Photoacoustic imaging of Cet-IRDye-AF conjugate containing phantoms. **(A)** Fluorescence, and **(C)** Photoacoustic imaging of increasing concentrations of Cet-IRDye-AF conjugate in tissue-mimicking phantoms. **(B)** Fluorescence imaging, and **(D)** Photoacoustic imaging of Cet-IRDye-AF with increasing degree of labeling of IRDye800, with and without trypsinization tissue-mimicking phantoms.

To confirm this, we further synthesized Cet-IRDye-AF with increasing IRDye800 payload (**Fig 1D**) and evaluated the effect of trypsinization on IRDye800 fluorescence and PA signal intensity. As shown in **Fig 6B**, IRDye800 fluorescence signal decreased with an increase in degree of labeling of IRDye800 on the antibody, possibly due to homoquenching at higher IRDye800 payloads. The fluorescence was however restored to a comparable value after trypsinization **(Fig 6B**). Interestingly, with increasing IRDye800 loading, the PA signal amplitude increased till a loading ratio of 1:3 (Cetuximab to IRDye800). The PA signal amplitude however decreased if the loading was further increased above 1:3 (**Fig 6D**). These results suggest that IRDye800 stacking as observed by a shoulder (at 703 nm) in the absorption spectra (**Fig 7B**) of the Cet-IRDye-AF conjugate contributes to an increased PA signal amplitude. This shoulder is characteristic of an H-aggregate observed with an increasing IRDye800 loading on the antibody (**Fig 7C**). While the H-aggregation appeared to increase even beyond a loading ratio of 1:3, the decrease in PA signal amplitude at a loading ratio of 1:5 was unexpected and could possibly be due to different modes of non-radiative relaxation of the excited state molecules.

**Figure 7:**
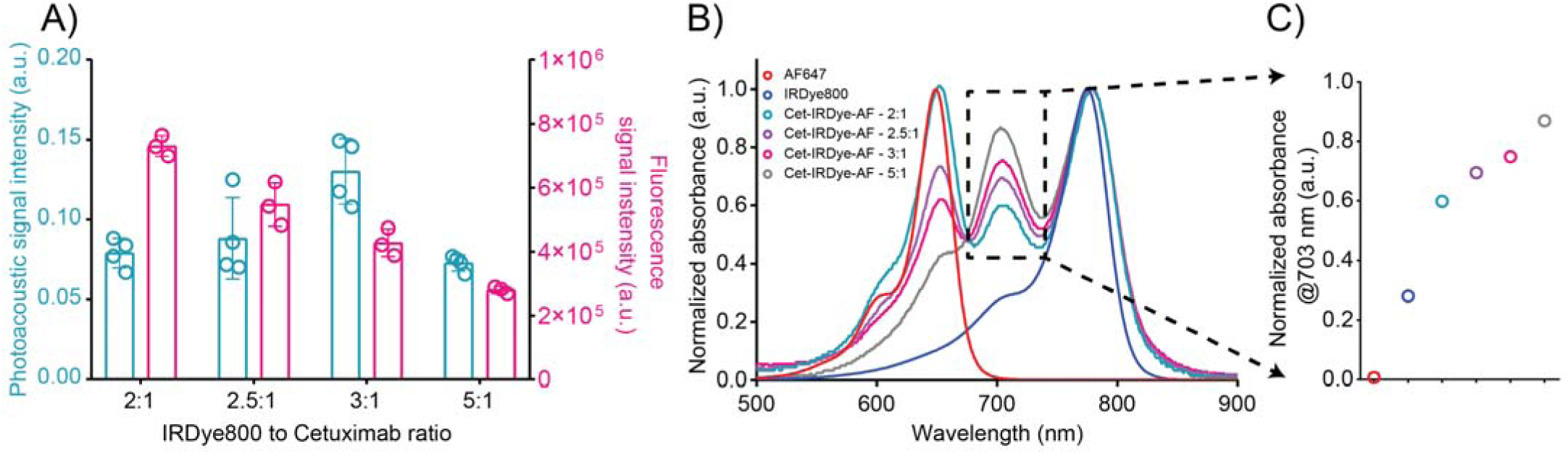
Characterization of spectral properties of H-aggregates formed by Cet-IRDye-AF conjugates. **(A)** Fluorescence imaging, and photoacoustic imaging of Cet-IRDye-AF with increasing degree of labeling of IRDye800 in tissue-mimicking phantoms (adapted from Fig 6B and 6D). **(B)** UV−vis spectra of Cet-IRDye-AF with increasing degree of labeling of IRDye800 in phosphate-buffered saline. **(C)** Absorption @703 nm, characteristic of H-aggregation for Cet-IRDye-AF with increasing degree of labeling of IRDye800.

## Discussion

While fluorescence-guided tumor surgeries have been in clinical practice for a few decades, there are certain limitations associated with the current practice of using fluorophores for guiding tumor resection surgeries. Two major limitations are the non-specific accumulation of fluorophores and the inherent optical limits of penetration (few mm) of NIR light used for fluorescence measurements. With recent advances in the development of therapeutic antibodies and antibody drug conjugates, several NIR fluorophore-antibody conjugates have been reported and are in clinical trials for detection and guiding tumor resection surgeries of which EGFR-targeted IRDye800 has been the most common with several clinical trials evaluating its efficacy in different tumor types.

This study discusses the potential of Cet-IRDye800 as a molecular-targeted photoacoustic contrast agent. While the utility of Cet-IRDye800 as a molecular-targeted fluorescence contrast agent is well established, its utility for PA imaging has been limited by several factors including low sensitivity for accurate detection. Nishio et al. demonstrated ex vivo PA imaging of lymph node metastasis using Pan-IRDye800 conjugates [13], however these results were not extendable to clinical studies possibly due to the lower light fluence permitted in the clinic and hence the lower detection limit of the imaging system [16]. More importantly, it has been identified that PA imaging is limited with its sensitivity and efforts have thus been made to develop novel agents with improved contrast for PA imaging [22-24].

In this study we highlight the importance of timing photoacoustic imaging post-injection of Cet-IRdye800 conjugates and possibly other similar antibody conjugates as well. We show that antibody conjugation promotes H-aggregation of IRDye800 which enhances PA signal intensity in time frames when the Cet-IRDye800 conjugate is intact. And, if imaging is delayed the PA signal intensity can decrease due to trypsinization of antibody-dye conjugate or degradation during the receptor mediated endocytosis (RME) pathway. This observation may have major implications in the way PA imaging is being performed in the clinic which thus far has yielded underwhelming results [16]. The study proposes a straight-forward strategy, that is shorter timeframes between infusion and imaging of molecular targeted PA contrast agents, to potentially capture significantly higher PA contrast using Cet-IRDye800 conjugates. We also propose an alternative to enhance this signal further through increasing the degree of labeling (DOL) to 3 instead of the clinically used degree of labeling of 2. PA imaging capability of Cet-IRDye800 with a DOL of 3 was however not tested in the current study but is expected to be higher than that obtained for Cet-IRDye with a DOL of 2, based on our in vitro studies. The importance of timing as proposed in this study is backed by other studies for different dyes - BHQ3 [9] and ICG [14]. For BHQ3-fluorescein trastuzumab conjugates imaging at 8 h post-injection yielded a higher PA signal in HER2 positive subcutaneous H2N tumors. The observed decrease in PA signal after the 8 h time-point was attributed to the metabolization of BHQ3 as has been demonstrated in earlier studies [29]. We assume the metabolization of BHQ3 is possibly following receptor mediated endocytosis of the BHQ3-fluorescein trastuzumab conjugate as it is observed in the time frames of the RME-based Cet-IRDye-AF degradation observed in our study. In the study by Wilson et al. [14], the RME of antibody-ICG conjugate was used to an advantage where the shift in the ICG absorption spectrum, and hence the spectroscopic PA signal, during cellular endocytosis and subsequent degradation of the B7-H3-ICG conjugate was exploited to differentiate between free B7-H3-ICG, non-specific controls, and B7-H3-ICG bound and internalized to its molecular target [14]. However, unlike ICG which shows a shift in absorption peak depending on its antibody bound or unbound status, IRDye800 does show an appearance of a shoulder peak at around 700 nm. This shoulder peak, however, is relatively low as compared to the primary absorption peak at 780 nm (at the DOLs tried in this study) which does not allow its utilization in PA imaging – 1) due to the lower absorption and 2) lower wavelength limiting its depth of penetration.

While the study demonstrates a possible methodology for capturing higher PA signals from Cet-IRDye800, currently being evaluated in the clinic for fluorescence imaging, there were a couple of limitations that must be addressed before this could be taken further. Firstly, the accuracy of PAI-based detection at such early time frames is a concern primarily due to the possible presence of unbound Cet-IRDye800. In microscopic evaluation of overlying skin and adjacent muscle tissue we did not observe Cet-IRDye-AF conjugate signal. However, macroscopic imaging of IRDye800 fluorescence did suggest an improvement in the tumor to background ratio from 6 h to 24 h time-point indicating the presence of Cet-IRDye-AF in relatively low quantities in the muscle and skin tissue at the 6 h timepoint. These low amounts may however be below the detection limit of PA imaging. Secondly, the distribution of Cet-IRDye-AF as observed in microscopic imaging was found to be significantly lower at the 6 h time-point as compared to the 24 h time-point. We compared Cet-IRDye-AF distribution between the two time-points by monitoring the Cet-IRDye-AF coverage area in the tumor cross-sections, the Cet-IRDye-AF to EGFR ratio to indicate the number of EGFR sites (as revealed by ex vivo EGFR staining) labeled by Cet-IRDye-AF, distance of Cet-IRDye-AF from the nearest CD31 positive blood vessel, and the Cet-IRDye-AF to CD31 ratio. All these parameters showed a significantly lower value at the 6 h timepoint as compared to the 24 h timepoint, suggesting the Cet-IRDye-AF was able to reach more EGFR positive tumor cells at the 24 h timepoint as compared to the 6 h timepoint. While this is not surprising, data from macroscopic imaging of IRDye800 suggested that IRDye800 fluorescence at the 24 h and 6 h were not significantly different. However, macroscopic imaging of AF647 showed a significant increase in fluorescence intensity with time due to unquenching. Since the *ex vivo* distribution studies were based on AF647 imaging which may have been quenched for the 6 h tumor samples, the differences between the different microscopic Cet-IRDye-AF distribution parameters may not be as significant as was observed in this study. Furthermore, the susceptibility of cyanine dyes including Indocyanine green (ICG) and IRDye800 to photobleaching under nanosecond laser illumination for PA imaging is another critical factor. In fact, Duffy et al. have demonstrated an ∼50% photobleaching in IRDye800 when exposed to 30 minutes of continuous PA imaging [30]. When performing 3D imaging of tumors across multiple wavelengths, the extended PA laser exposure exacerbates photobleaching effects. This could lead to underestimations of dye accumulation in the tumors and could potentially skew quantitative results compared to fluorescence images from AF647. Third, the tumors studied in this study were fairly big, close to ∼ 1 cm and therefore had a larger Cet-IRDye-AF accumulation and hence high enough PA signal that could be detected. Future studies should focus on identifying lymph node metastasis using the Cet-IRDye-AF conjugate and whether those could be efficiently detected using PA imaging at early time-points.

While with this study we attempt to emphasize the importance of timing for capturing a high PA signal intensity. We believe this could be further improved through one of the following three strategies – 1) As discussed earlier increasing the IRDye800 DOL to 3 can possibly enhance PA signal amplitude. 2) The high receptor expression in A431 tumors used here may have posed a significant binding-site-barrier limiting the extent of migration, distribution and accumulation of Cet-IRDye-AF conjugate thus decreasing the PA signal. Pre-injecting animals with free Cetuximab can possibly help in improving the PA signal amplitude [31]. 3) Performing photodynamic priming (PDP) prior to Cet-IRDye-AF conjugate administration can help in increased accumulation of the conjugate at relatively early timepoints thereby enhancing the tumor accumulation and hence PA signal intensity [32]. This has been exemplified in pre-clinical oral cancer models where PDP at 75 J/cm^2^ expedited interstitial accumulation of Cet-IRDye800 by 10.5-fold and increased its fractional tumor coverage by 49.5%, 1 h after administration [33].

In summary, this study demonstrates the importance of timing in current PA imaging regimens using Cet-IRDye800 and provides mechanistic evidence of why performing PA imaging at early timepoints can lead to a better PA signal quantity. In addition, we propose several strategies to enhance the PA signal further. We believe the current study can prove to be transformative in the way PA imaging is being performed in the clinic, specifically for lymph node imaging for identifying occult metastasis.

## Acknowledgements

This work was supported by grants from National Institute of Health R01 CA266701 to SM, and P01 CA084203, R01 CA231606, and S10OD012326 to TH.

## Author Contributions

Conceptualization, M.A.S., S.M., and T.H.; methodology, M.A.S., D.A., A.S., M.X., and S.M.; formal analysis, M.A.S., D.A., A.S., M.X., and S.M.; resources, S.M., and T.H; data curation, M.A.S., D.A., and M.X.; writing— original draft preparation, M.A.S., D.A., and S.M.; writing—review and editing, M.A.S., D.A., A.S., M.X., S.M., and T.H.; supervision, T.H.; project administration, T.H.; funding acquisition, T.H., and S.M.

## References

1. Wang K, Du Y, Zhang Z, He K, Cheng Z, Yin L, et al. Fluorescence image-guided tumour surgery. Nature Reviews Bioengineering. 2023; 1: 161–79.

2. Troyan SL, Kianzad V, Gibbs-Strauss SL, Gioux S, Matsui A, Oketokoun R, et al. The FLARE™ intraoperative near-infrared fluorescence imaging system: a first-in-human clinical trial in breast cancer sentinel lymph node mapping. Annals of surgical oncology. 2009; 16: 2943–52.

3. Stummer W, Novotny A, Stepp H, Goetz C, Bise K, Reulen HJ. Fluorescence-guided resection of glioblastoma multiforme utilizing 5-ALA-induced porphyrins: a prospective study in 52 consecutive patients. Journal of neurosurgery. 2000; 93: 1003–13.

4. Ferroli P, Acerbi F, Albanese E, Tringali G, Broggi M, Franzini A, et al. Application of intraoperative indocyanine green angiography for CNS tumors: results on the first 100 cases. Intraoperative Imaging: Springer; 2011. p. 251-7.

5. Obaid G, Spring BQ, Bano S, Hasan T. Activatable clinical fluorophore-quencher antibody pairs as dual molecular probes for the enhanced specificity of image-guided surgery. Journal of biomedical optics. 2017; 22: 121607.

6. Spring BQ, Abu-Yousif AO, Palanisami A, Rizvi I, Zheng X, Mai Z, et al. Selective treatment and monitoring of disseminated cancer micrometastases in vivo using dual-function, activatable immunoconjugates. Proceedings of the National Academy of Sciences. 2014; 111: E933–E42.

7. Folli S, Wagnieres G, Pelegrin A, Calmes J, Braichotte D, Buchegger F, et al. Immunophotodiagnosis of colon carcinomas in patients injected with fluoresceinated chimeric antibodies against carcinoembryonic antigen. Proceedings of the National Academy of Sciences. 1992; 89: 7973–7.

8. Rosenthal EL, Warram JM, De Boer E, Chung TK, Korb ML, Brandwein-Gensler M, et al. Safety and tumor specificity of cetuximab-IRDye800 for surgical navigation in head and neck cancer. Clinical Cancer Research. 2015; 21: 3658–66.

9. Maeda A, Bu J, Chen J, Zheng G, DaCosta RS. Dual in vivo photoacoustic and fluorescence imaging of HER2 expression in breast tumors for diagnosis, margin assessment, and surgical guidance. Mol Imaging. 2015; 14: 7290.2014. 00043.

10. Park S, Kim J, Jeon M, Song J, Kim C. In vivo photoacoustic and fluorescence cystography using clinically relevant dual modal indocyanine green. Sensors. 2014; 14: 19660–8.

11. Zhang D, Zhao Y-X, Qiao Z-Y, Mayerholffer U, Spenst P, Li X-J, et al. Nano-confined squaraine dye assemblies: new photoacoustic and near-infrared fluorescence dual-modular imaging probes in vivo. Bioconjugate chemistry. 2014; 25: 2021–9.

12. Park J, Choi S, Knieling F, Clingman B, Bohndiek S, Wang LV, et al. Clinical translation of photoacoustic imaging. Nature Reviews Bioengineering. 2024.

13. Nishio N, van den Berg NS, Martin BA, van Keulen S, Fakurnejad S, Rosenthal EL, et al. Photoacoustic Molecular Imaging for the Identification of Lymph Node Metastasis in Head and Neck Cancer Using an Anti-EGFR Antibody–Dye Conjugate. Journal of Nuclear Medicine. 2021; 62: 648–55.

14. Wilson KE, Bachawal SV, Abou-Elkacem L, Jensen K, Machtaler S, Tian L, et al. Spectroscopic Photoacoustic Molecular Imaging of Breast Cancer using a B7-H3-targeted ICG Contrast Agent. Theranostics. 2017; 7: 1463–76.

15. Becker C, Hardarson J, Hoelzer A, Geisler A, Schulz T, Reichl C, et al. Evaluation of cervical lymph nodes using multispectral optoacoustic tomography: a proof-of-concept study. European Archives of Oto-Rhino-Laryngology. 2023; 280: 4657–64.

16. Vonk J, Kukačka J, Steinkamp PJ, de Wit JG, Voskuil FJ, Hooghiemstra WTR, et al. Multispectral optoacoustic tomography for in vivo detection of lymph node metastases in oral cancer patients using an EGFR-targeted contrast agent and intrinsic tissue contrast: A proof-of-concept study. Photoacoustics. 2022; 26: 100362.

17. Te Velde E, Veerman T, Subramaniam V, Ruers T. The use of fluorescent dyes and probes in surgical oncology. European Journal of Surgical Oncology. 2010; 36: 6–15.

18. Zhang Y, Jeon M, Rich LJ, Hong H, Geng J, Zhang Y, et al. Non-invasive multimodal functional imaging of the intestine with frozen micellar naphthalocyanines. Nature Nanotechnology. 2014; 9: 631–8.

19. Wood CA, Han S, Kim CS, Wen Y, Sampaio DRT, Harris JT, et al. Clinically translatable quantitative molecular photoacoustic imaging with liposome-encapsulated ICG J-aggregates. Nature Communications. 2021; 12: 5410.

20. Tzoumas S, Ntziachristos V. Spectral unmixing techniques for optoacoustic imaging of tissue pathophysiology. Philos Trans A Math Phys Eng Sci. 2017; 375: 20170262.

21. A. PS. Optical Absorption of Hemoglobin. https://omlcorg/spectra/hemoglobin/. 1999.

22. Saad MA, Grimaldo-Garcia S, Sweeney A, Mallidi S, Hasan T. Dual-Function Antibody Conjugate-Enabled Photoimmunotherapy Complements Fluorescence and Photoacoustic Imaging of Head and Neck Cancer Spheroids. Bioconjugate Chemistry. 2024; 35: 51–63.

23. Saad MA, Pawle R, Selfridge S, Contreras L, Xavierselvan M, Nguyen CD, et al. Optimizing Axial and Peripheral Substitutions in Si-Centered Naphthalocyanine Dyes for Enhancing Aqueous Solubility and Photoacoustic Signal Intensity. Int J Mol Sci. 2023; 24: 2241.

24. Saad MA, Xavierselvan M, Sharif HA, Selfridge S, Pawle R, Varvares M, et al. Dual Function Antibody Conjugates for Multimodal Imaging and Photoimmunotherapy of Cancer Cells. Photochem Photobiol. 2022; 98: 220–31.

25. Saad MA, Zhung W, Stanley ME, Formica S, Grimaldo-Garcia S, Obaid G, et al. Photoimmunotherapy Retains Its Anti-Tumor Efficacy with Increasing Stromal Content in Heterotypic Pancreatic Cancer Spheroids. Molecular Pharmaceutics. 2022; 19: 2549–63.

26. Zinn KR, Korb M, Samuel S, Warram JM, Dion D, Killingsworth C, et al. IND-Directed Safety and Biodistribution Study of Intravenously Injected Cetuximab-IRDye800 in Cynomolgus Macaques. Molecular Imaging and Biology. 2015; 17: 49–57.

27. Hynes NE, Lane HA. ERBB receptors and cancer: the complexity of targeted inhibitors. Nature Reviews Cancer. 2005; 5: 341–54.

28. Lu G, Fakurnejad S, Martin BA, van den Berg NS, van Keulen S, Nishio N, et al. Predicting Therapeutic Antibody Delivery into Human Head and Neck Cancers. Clinical Cancer Research. 2020; 26: 2582–94.

29. Linder KE, Metcalfe E, Nanjappan P, Arunachalam T, Ramos K, Skedzielewski TM, et al. Synthesis, In Vitro Evaluation, and In Vivo Metabolism of Fluor/Quencher Compounds Containing IRDye 800CW and Black Hole Quencher-3 (BHQ-3). Bioconjugate Chemistry. 2011; 22: 1287-97.

30. Duffy MJ, Planas O, Faust A, Vogl T, Hermann S, Schäfers M, et al. Towards optimized naphthalocyanines as sonochromes for photoacoustic imaging in vivo. Photoacoustics. 2018; 9: 49–61.

31. Lu G, Nishio N, van den Berg NS, Martin BA, Fakurnejad S, van Keulen S, et al. Co-administered antibody improves penetration of antibody–dye conjugate into human cancers with implications for antibody–drug conjugates. Nature Communications. 2020; 11: 5667.

32. Bhandari C, Fakhry J, Eroy M, Song JJ, Samkoe K, Hasan T, et al. Towards Photodynamic Image-Guided Surgery of Head and Neck Tumors: Photodynamic Priming Improves Delivery and Diagnostic Accuracy of Cetuximab-IRDye800CW. Front Oncol. 2022; 12.

33. Nguyen A, Bhandari C, Keown M, Malkoochi A, Quaye M, Mahmoud D, et al. Increasing the Dye Payload of Cetuximab-IRDye800CW Enables Photodynamic Therapy. Molecular Pharmaceutics. 2024; 21: 3296–309.

