## Supplementary fig S1, S2 and S3 for "Temporal dynamics of fluorescence and photoacoustic signals of a Cetuximab-IRDye800 conjugate in EGFR-overexpressing tumors"

Tayyaba Hasan Ph.D.

Professor of Dermatology

Professor of Health Sciences and Technology (Harvard-MIT)

Wellman Center for Photomedicine, Harvard Medical School

Massachusetts General Hospital

40 Blossom Street, (Bartlett 314), Boston, MA 02114

**Key words:** Photoacoustic imaging, fluorescence imaging, image-guided therapy, molecular-targeted imaging probes, photodiagnosis, optical imaging

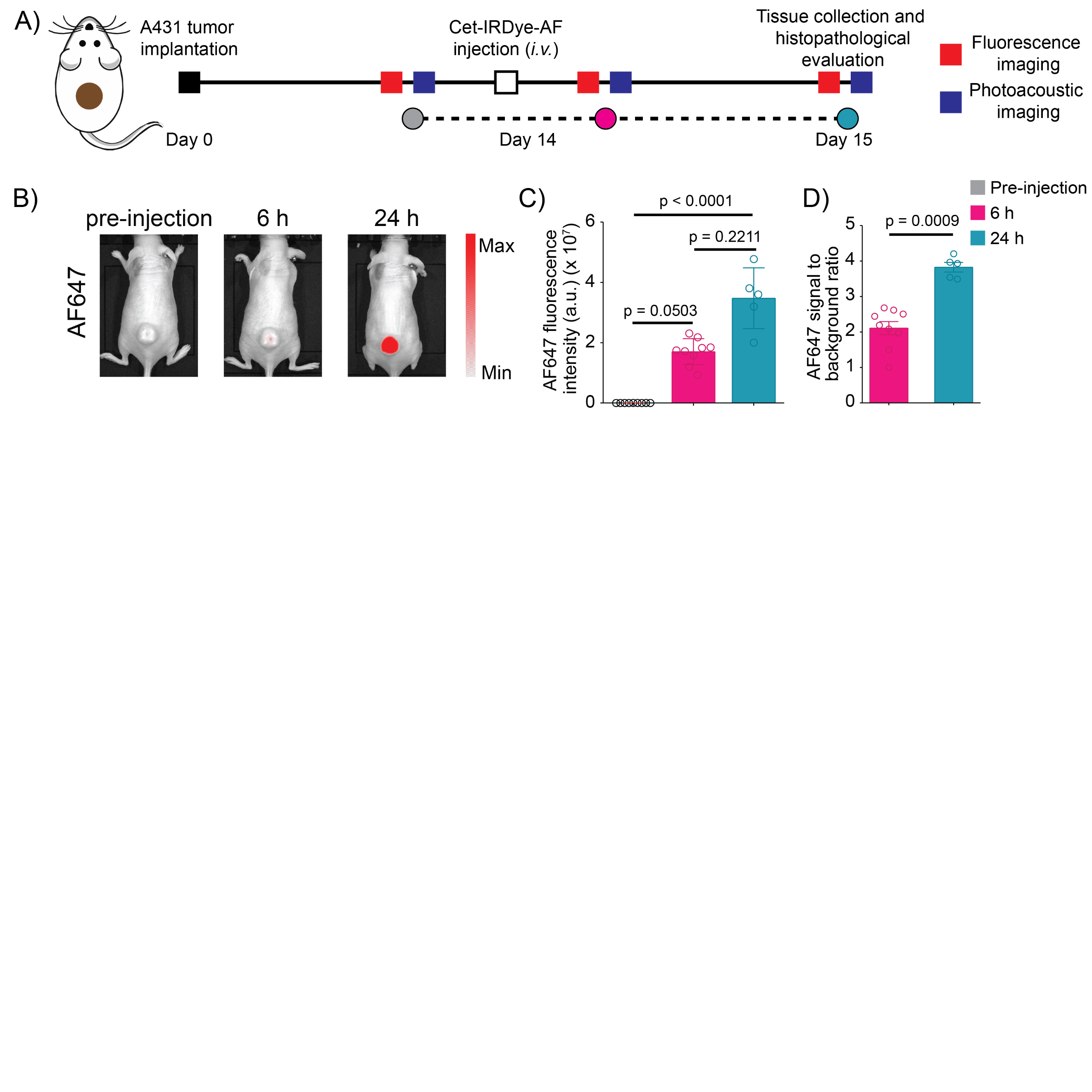

**Supplementary figure 1: Temporal dynamics of AF647 fluorescence signals of Cet-IRDye-AF (A)** Experimental timeline of in vivo dual modal imaging in A431 subcutaneous xenograft model. **(B)** In vivo fluorescence imaging of tumor bearing mice before injection and 6 h, and 24 h after injection of Cet-IRDye-AF conjugate. **(C)** Quantification of absolute AF647 fluorescence signal intensity at different time-points before and after Cet-IRDye-AF injection. **(D)** Tumor to background fluorescence ratio at different time-points before and after Cet-IRDye-AF injection. Data are presented as mean ± SD (n ≥ 5 per group), analyzed using Kruskal-Wallis test, followed by post-hoc pairwise comparisons using Dunn's test. P-values are provided for each graph.

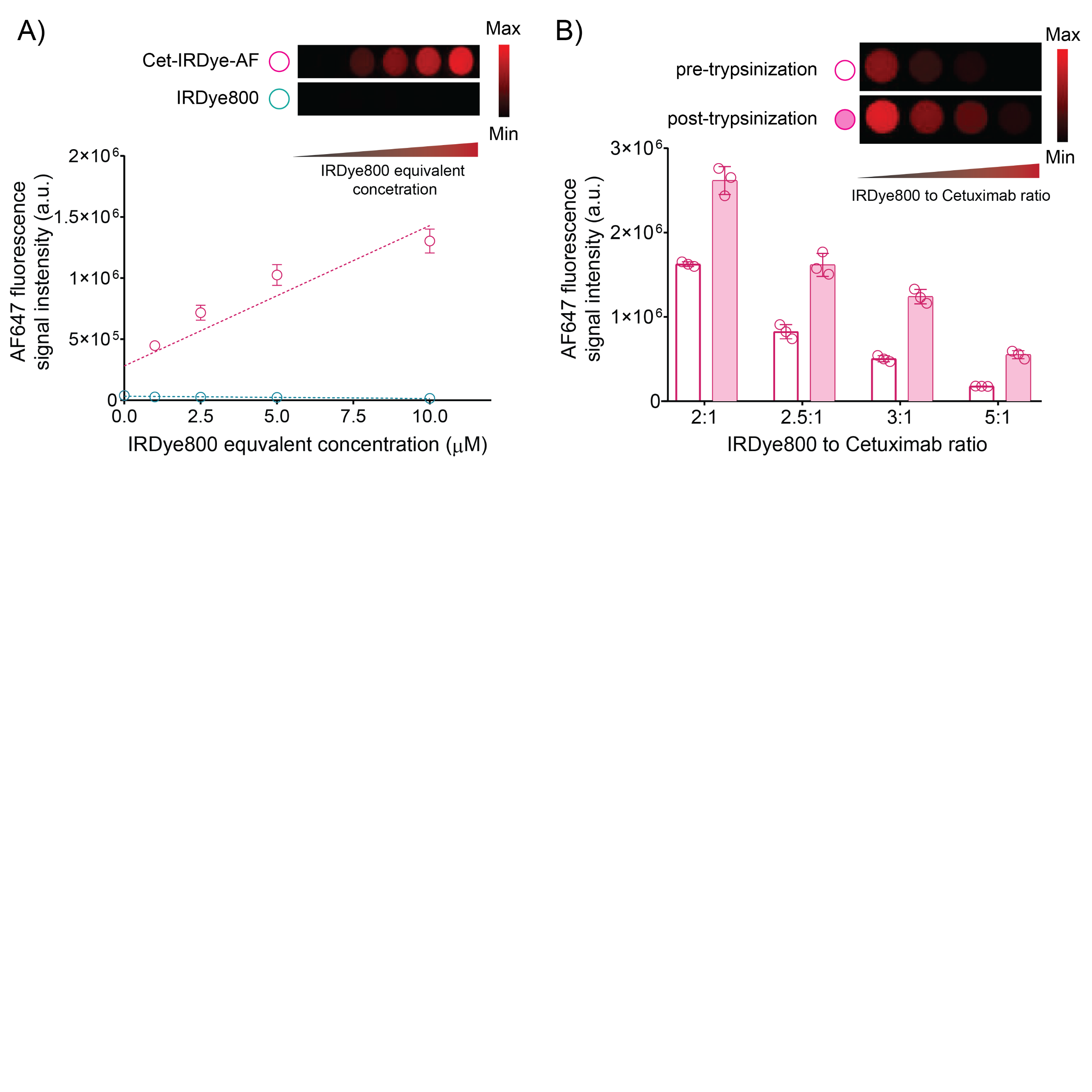

**Supplementary figure 2: AF647 fluorescence imaging of Cet-IRDye-AF conjugate containing phantoms. (A)** Fluorescence imaging of phantoms prepared with increasing concentrations of Cet-IRDye-AF conjugate in tissue-mimicking phantoms. **(B)** Fluorescence imaging of Cet-IRDye-AF with increasing degree of labeling of IRDye800, with and without trypsinization tissue-mimicking phantoms.

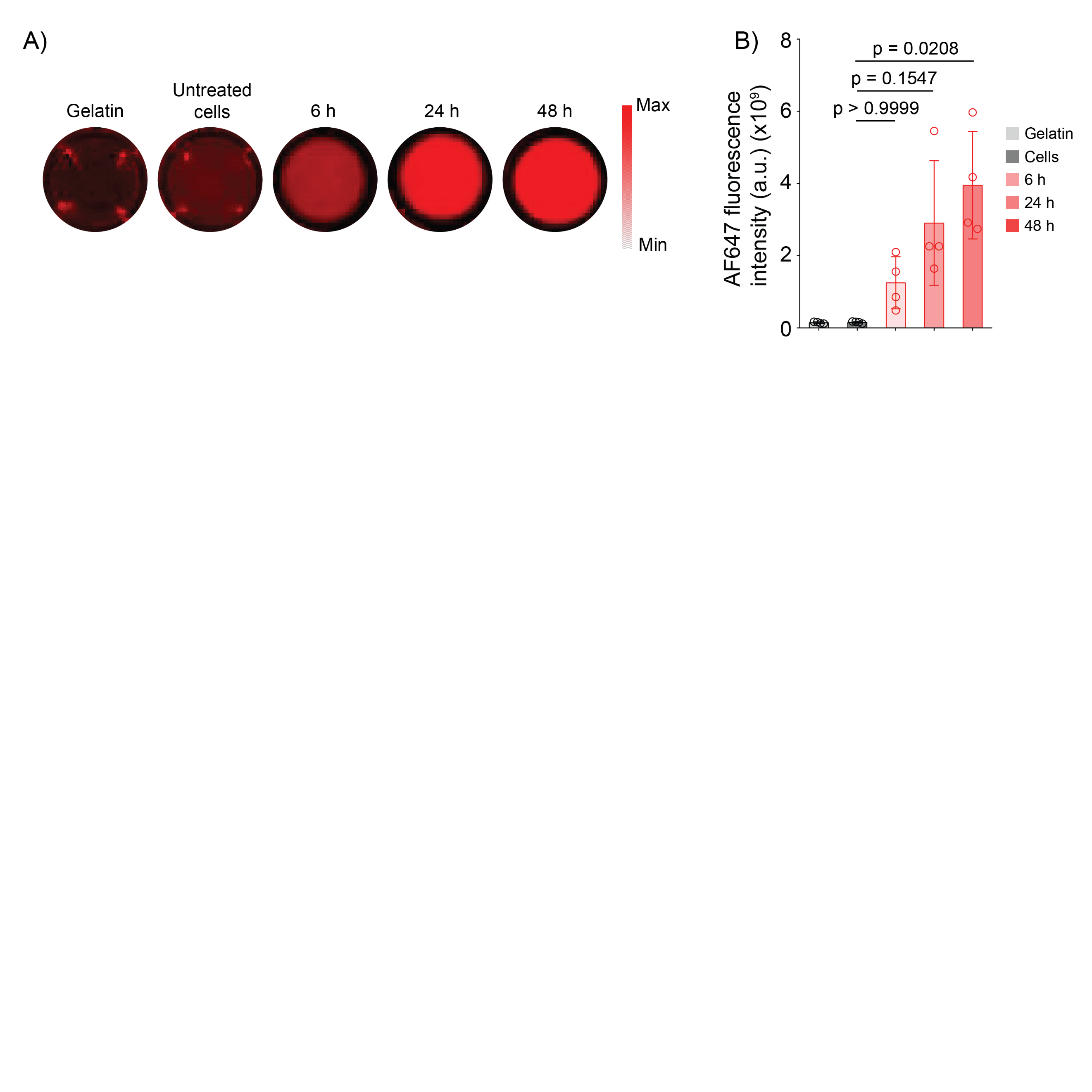

**Supplementary figure 2: (A)** IVIS images of AF647 fluorescence signal from Cet-IRDye-AF-treated A431 cells prepared as gelatin embedded tissue mimicking phantoms in a 96-well plate. **(B)** Quantification of fluorescence signals from the IVIS images. Incubation of A431 cells with Cet-IRDye-AF showed a steady increase in AF647 fluorescence signals with time. Data are presented as mean ± SD (n = 4 tumor mimicking phantoms per group for fluorescence images. Data is analyzed using Kruskal-Wallis test, followed by post-hoc pairwise comparisons using Dunn's test. P-values are provided for each graph.
